# Covalent Solvatochromic Membrane Probe for Sensing Lipid Surrounding of Proteins

**DOI:** 10.64898/2026.10.03.756383

**Authors:** Martijn Visser, Tiffany Loo Chin Moy, Rémi Pelletier, Andrey S. Klymchenko

## Abstract

Plasma membranes (PM) have been traditionally studied by fluorescent probes by either labelling lipid or protein parts. Based on our MemGraft concept, we report a fluorescent probe based on solvatochromic dye Nile Red (MemGraft-NR) for sensing lipid organization in the immediate surrounding of proteins, i.e. at the protein-lipid interface. Two sequentially attached MemGraft scaffolds are required to ensure robust grafting of Nile Red to PM. In contrast to classical lipid-bound membrane probes, MemGraft-NR exhibits high resistance to the presence of serum and fixation with permeabilization. Two-color confocal imaging of MemGraft-NR in live cells reveals higher polarity at the protein-lipid interface compared to lipid-bound probe NR12A. Surprisingly, MemGraft-NR displays higher sensitivity to variation of cholesterol content in PM compared to NR12A, suggesting that protein-lipid interface is particularly impacted by changes in the lipid composition. MemGraft-NR also enabled in situ monitoring of lipid exchange and permeabilization of PM, something impossible for classical solvatochromic PM probes. Unexpectedly, MemGraft-NR reveals a dramatic decrease in the local polarity in early endosomes, suggesting that endosomal negative curvature provides strong impact on organization of the protein-lipid interface. The developed probe opens a new way to study biological membranes by sensing lipid surrounding of membrane proteins.

## Introduction

The cellular plasma membrane (PM) is a complex multifunctional platform composed of a great variety of phospholipids, sterols, proteins, polysaccharides and other molecules and presenting complex lateral and transmembrane organization.^1-3^ The unmet need to understand better the PM stimulated the development of multiple chemical tools, including fluorescent probes.^4-5^ A vast majority of these probes are based on non-covalent interactions, such as lipophilic or amphiphilic probes,^6-12^ fluorescent antibodies^13^ and lectin-bound probes.^14^ However, the non-covalent strategies, despite their simplicity of use, bear a number of drawbacks: (a) the loss of labelling by desorption to biological serum-containing medium; (b) exchange with non-labelled cells; (c) incompatibility with fixation, especially permeabilization, tissue clearance or expansion microscopy protocols.

Covalent approaches include fusion of membrane proteins to exogenous proteins such as fluorescent proteins,^15^ the use of protein tags,^16-19^ or introduction of unnatural amino acids^20^ which require genetic modification and rather long sample preparation times (∼24h). Chemical approaches of cell surface labelling are rather rare, as they require a design of reagents providing covalent chemistry specifically at the cell surface. Here, one should mention metabolic labelling, in particular using biorthogonal click chemistry.^21-22^ However, this method also requires relatively long sample preparation related to metabolic insertion of the reactive groups. Other methods exploit ligand-directed protein labelling, where ligand targeting triggers covalent labelling.^23-24^ However, those methods require the presence of a specific receptor of the cell surface and the labelling density is limited by the expression of the target receptor. Recently we introduced the MemGraft family of probes (Figure 1A), which operates by lipid-directed covalent labelling of the cell surface. They use a low affinity membrane anchor to target the cell membrane that facilitates grafting to membrane proteins through active ester chemistry in just 5 min.^25^ These probes overcame limitations of their non-covalent analogues, making them compatible with serum-containing media, fixation, permeabilization, trypsinization, co-culture of differently labelled cells, long-term imaging, barcoding and cell manipulation. However, so far this MemGraft strategy was used exclusively for cell labelling, without a sensing function. We consider that lipid-directed protein labelling could be a unique opportunity to study lipid surrounding of membrane proteins,^26^ which is currently very difficult to address with conventional probes. As MemGraft strategy targets any membrane protein, such a probe would sense the properties of protein-lipid interface in plasma membranes. A powerful method to study molecular environment, its organization and biomolecular interactions is to use so-called environment-sensitive dyes.^27^ They sense local properties of their close surrounding by changing their fluorescence intensity or color. The key representatives include solvatochromic dyes sensitive to polarity,^27-28^ molecular rotors sensitive to viscosity,^29-30^ flippers sensitive to the lipid order and membrane tension,^9,31^ etc. Solvatochromism is the ability of a fluorophore to change its fluorescence spectrum depending on the solvent polarity.^12,13^ The more polar environments allow for stabilization of the excited state of the fluorophore, thus lowering the energy and shifting the fluorescence emission towards longer wavelengths (red shift). This property has been extensively used to study organization of lipid membranes and monitor a variety of biomolecular interactions, including peptide-membrane, protein-protein, and protein-DNA interactions.^4,27^ Solvatochromic dyes were particularly powerful in the applications to study lipid organization in model and cell membranes.^32-33^ Thus, solvatochromic dyes such as Laurdan,^34^ styryl dyes,^35^ push-pull pyrene,^36^ and Nile Red and its derivatives (e.g. NR12A, Figure 1A)^6-7^ have previously been described to change their fluorescence depending on the lipid order in model and cell membranes. As a general rule, decrease in the lipid order increases local polarity and hydration of the lipid membrane, thus leading to observed red shifts in the emission of solvatochromic dyes. ^32-33^ We considered that the extension of the MemGraft concept to a solvatochromic dye would enable sensing lipid environment of the cells in the proximity of membrane proteins, which provide a completely new view on the lipid organization as compared to “bulk” lipid order of biomembranes.

**Figure 1.**
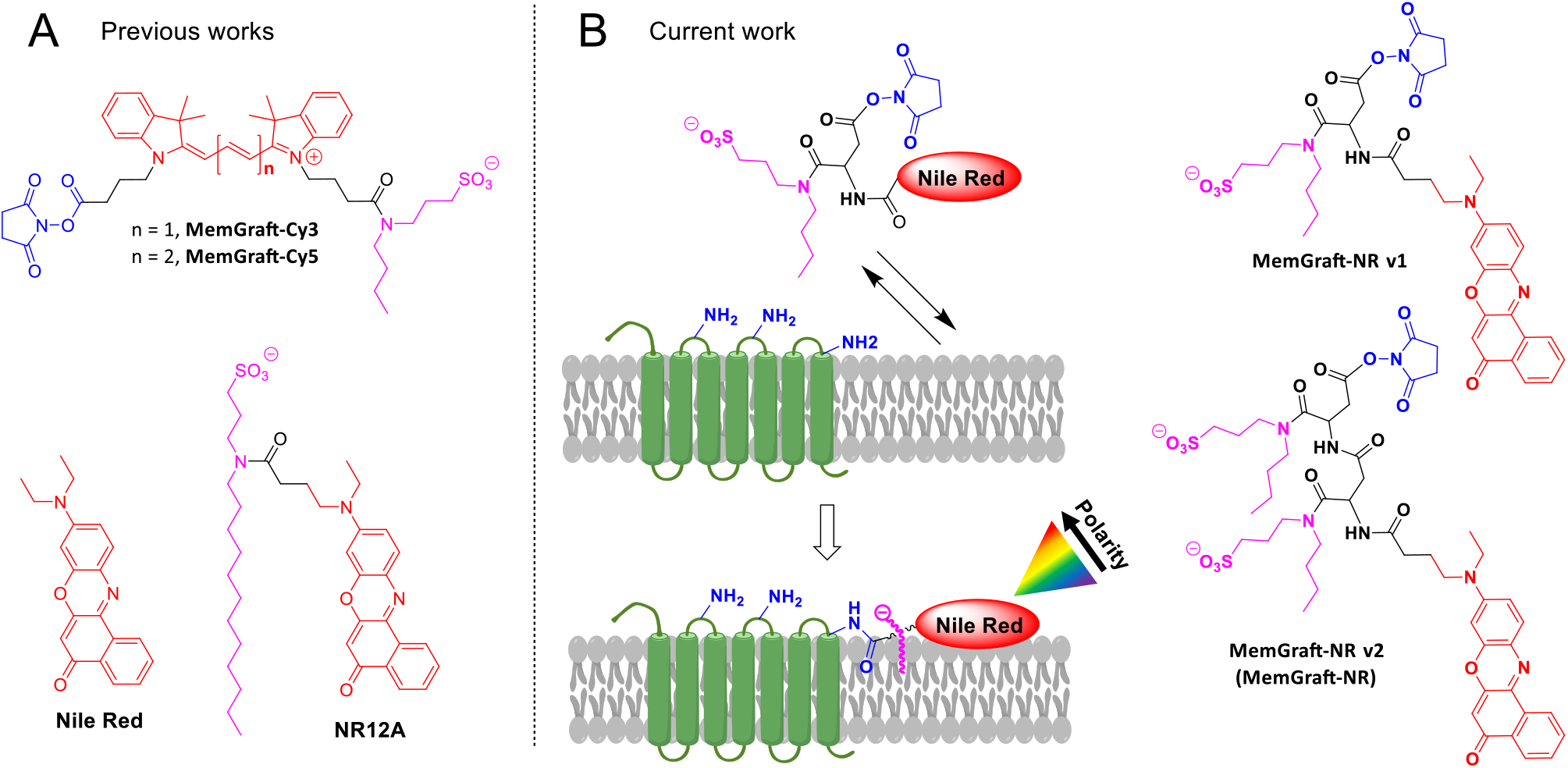
Principle of MemGraft and solvatochromic membrane probes. (A) Chemical structure of previously developed covalent MemGraft probes and non-noncovalent solvatochromic probe NR12A based on Nile Red and (B) current work: scheme of MemGraft labelling with solvatochromic dye Nile Red and two version of MemGraft probes synthesized in this work.

In this work, we report a polarity-sensitive MemGraft probe based on Nile Red (MemGraft-NR) fluorophore that enables sensing lipid surrounding of membrane proteins. To this end we developed a MemGraft scaffold containing both a lipid anchor and NHS-ester functions. We found that the single MemGraft scaffold was insufficient due to rapid internalization, whereas a double scaffold yielded an efficient PM labelling probe. Owing to the covalent grafting to the PM, the new probe showed resistance to serum-containing medium and to permeabilization, which washed away a reference non-covalent reference dye based on Nile Red (NR12A). MemGraft-NR exhibited high sensitivity to changes in the lipid composition, which surpassed even the lipid-bound probe NR12A. The probe also allowed for monitoring of lipid exchange, cell fixation and lipid removal during permeabilization, something difficult to realize with classical probes. Finally, we showed that MemGraft-NR is a useful tool to study endocytosis, where it revealed critical changes at the protein-lipid interface during the endosome formation. Overall, we introduce a tool, that allows a different look at the biomembrane by focusing on environment properties at the protein-lipid interface, notably the organization of the lipid surroundings of membrane proteins.

## Results and discussion

### Probe design

In contrast to cyanine dyes previously used in the design of MemGraft probes,^25^ the NR-COOH Nile Red fluorophore is monofunctional. This prompted us to develop a MemGraft scaffold **3**, which can adapt to any monofunctional dye using a protected aspartic acid and 3-(butylamino)propane-1-sulfonate **1** (Scheme 1). The latter was previously found to ensure reversible low-affinity binding of Nile Red to the cell membrane surface without cell permeation,^6^ while the former would provide a carboxylate function required for grafting to the membrane proteins (Figure 1B).^25^ Since the Nile Red core is neutral and the cyanines are charged, we hypothesized that the Nile Red probe will not behave the same way. Thus, two versions of the Nile Red MemGraft (MemGraft-NR or MGNR) probe were synthesized (i) with only one universal anchor (MemGraft-NR v1), and (ii) with a double anchor (MemGraft-NR v1, Figure 1B). To this end, the MemGraft scaffold was first obtained by amide coupling t-butyl and Fmoc-protected aspartic acid with anchor **1** followed by Fmoc removal (Scheme 1). Next, NR-COOH was coupled to the MemGraft scaffoldand the t-butyl was removed to yield non-activated MemGraft **4**. Finally, the latter was activated to the NHS ester, which corresponds to MemGraft-NR v1 (Scheme 1). To obtain MemGraft-NR v2 version, compound **4** was coupled again with MemGraft scaffold **3**, followed by tert-butyl group removal and carboxylate activation to the NHS ester (Scheme 1).

**Scheme 1.**
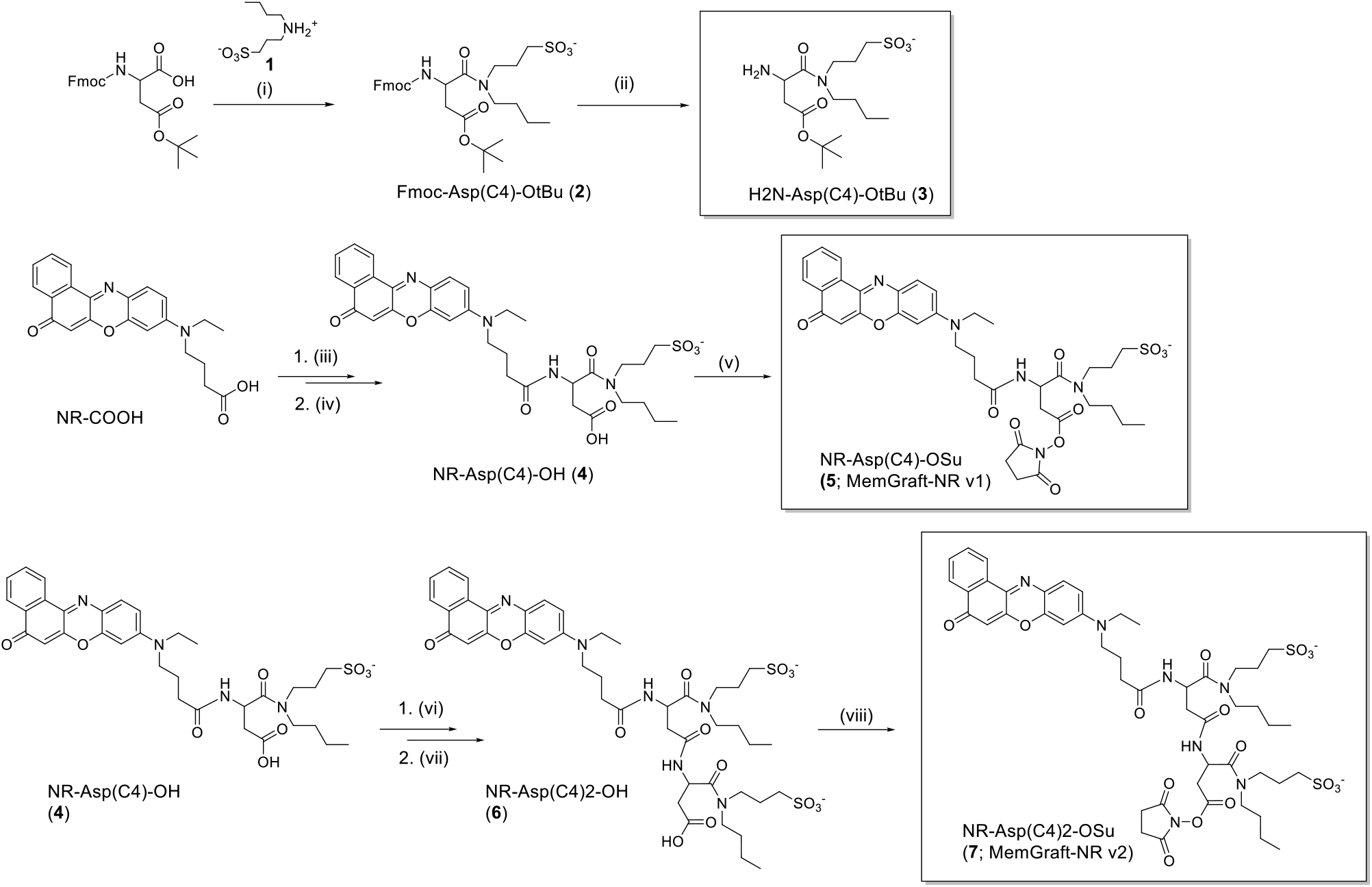
Synthesis of MemGraft scaffold **3** and MemGraft-NR probes. (i) **1** (1.1 eq), PyBOP (1.1 eq), DIPEA (3.5 eq), DMF, argon, r.t., 18h, 51%. (ii) Piperidine/DMF (20% v/v), r.t., 30 mins, 68%. (iii) **3** (1.1 eq), PyBOP (1.1 eq), DIPEA (5 eq) DMF argon r.t. 18h, 50%. (iv) TFA/DCM (1:5) r.t. 1h, quant. (v) TSTU (1.5 eq), DIPEA (1.5 eq), DCM, r.t., 4h, 14% isolated yield. (vi) **3** (1.1 eq), PyBOP (2 eq), DMF, argon, r.t., 18h, 21%. (vii) TFA/DCM (1:5) r.t. 1h, quant. (viii) TSTU (2 eq), DIPEA (2 eq), DMF/ACN (1;1), r.t., 4h, quant.

### Membrane staining

In order to verify membrane staining, colocalization experiments with a reference membrane probe MemBright-640 (MB640)^8^ were performed. MB640 was used to prevent interference with the Nile Red excitation at 488 nm and emission from 550 to 650 nm.

Control experiments using non-activated probes were performed to exclude the possibility of non-covalent binding of the NR probes with the membrane. Staining with MemGraft-NR (MGNR) v1 and v2, as well as their non-activated analogues, was performed as previously described^25^ for only 5 minutes, with washing steps to remove excess of the dyes. The reference probe MemBright-640 was then added to the final imaging media.

Surprisingly, MemGraft-NR v1 with only one anchor and its non-activated control rapidly stained the interior of the cell without colocalization with MB640 at the PM. This suggests that Nile Red-based probe does not behave the same way as Cyanine-based MemGraft, where one anchor was sufficient for membrane staining.^25^ The activated probe MemGraft-NR v2 displayed good membrane staining and colocalization with MB640, as well as little intracellular signal (Figure 2). In contrast, the non-activated control of MemGraft-NR v2 showed no membrane labelling and very little internalization (Figure 2). We can conclude that MemGraft-NR v2 stains selectively the cell plasma membrane by a chemical grafting with its NHS ester. Moreover, a single anchor group was insufficient, so that the second anchor was found crucial for specific labelling. We can speculate that Nile Red fluorophore is more hydrophobic than cyanines and requires two sulphonate groups with two short chains to prevent internalization through the plasma membrane and minimize non-specific binding to the cell surface. Therefore, all further studies were focused on MemGraft-NR v2, which will be called MemGraft-NR for simplicity.

**Figure 2.**
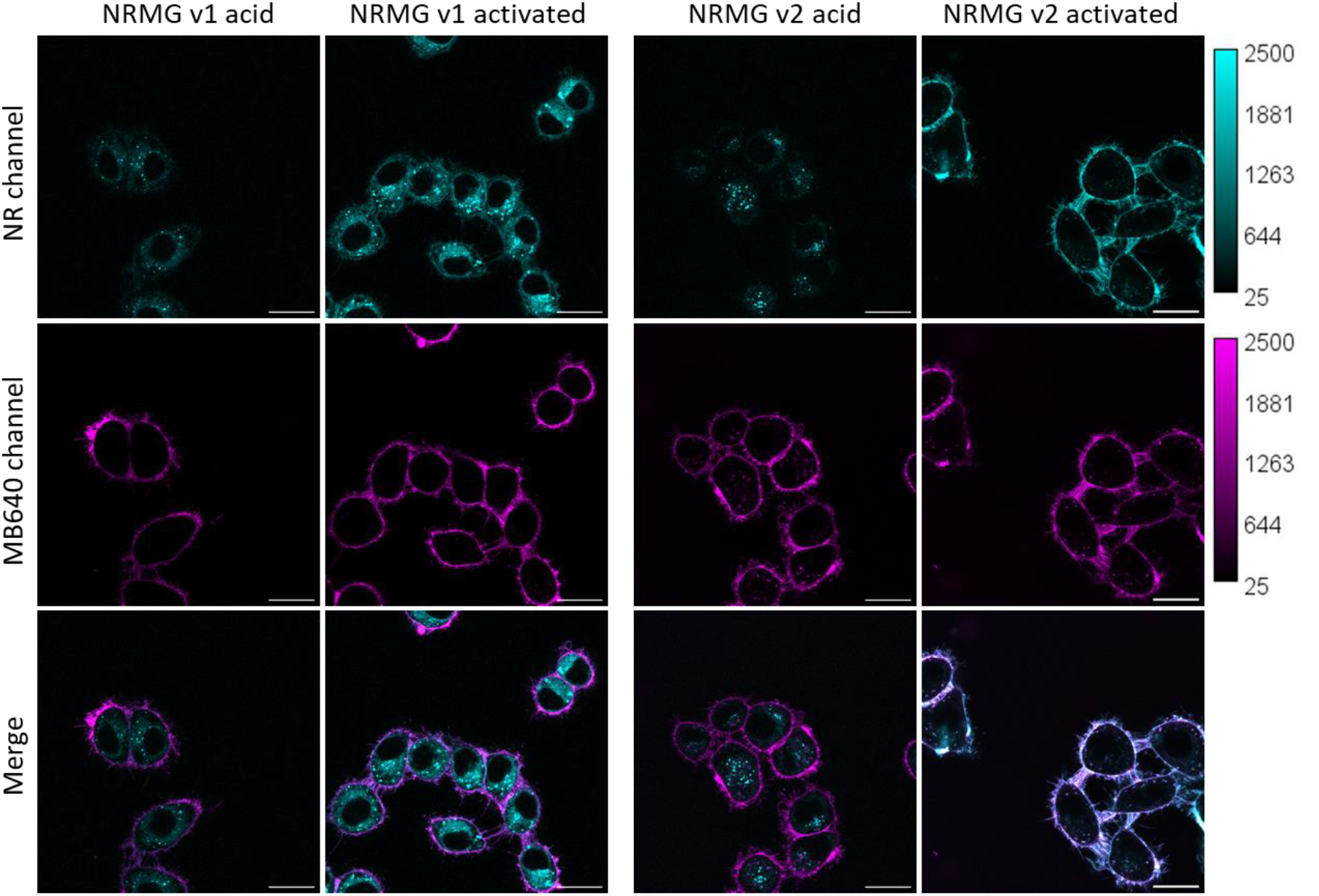
Confocal microscopy images of live KB cells in OptiMem stained with MemGraft-NR v1 and v2 probes or their non-activated analogues, and MemBright 640 as a reference membrane probe. MemGraft and MB640 dyes were excited using 488 and 638 nm laser, respectively. All images use the same intensity scale. Scale bars: 20 µm.

The advantage of MemGraft-type probes compared to “traditional” lipophilic membrane probes is the resistance to serum.^25^ As MemGraft probes are attached covalently to membrane proteins they cannot be extracted by serum albumins, in contrast to probes that rely on physical interaction with the membrane. To verify this for MemGraft-NR, we used previously reported membrane probe NR12A based on Nile Red fluorophore that binds non-covalently to the lipid membrane. In the serum containing medium, the fluorescence intensity of NR12A at the cell PM decreased over time, whereas MemGraft-NR signal remained on the PM for at least for 30 min tested, showing only a little decrease (Figure 3). A long-term imaging experiment was performed, where cells stained with MemGraft-NR and incubated in full serum containing medium were imaged at different time points. Over time, we can see that the signal intensity decreases as the cells divide and internalize their membranes, however, a signal still remains at the membrane even after 24 and 48 h (Figure S1).

**Figure 3.**
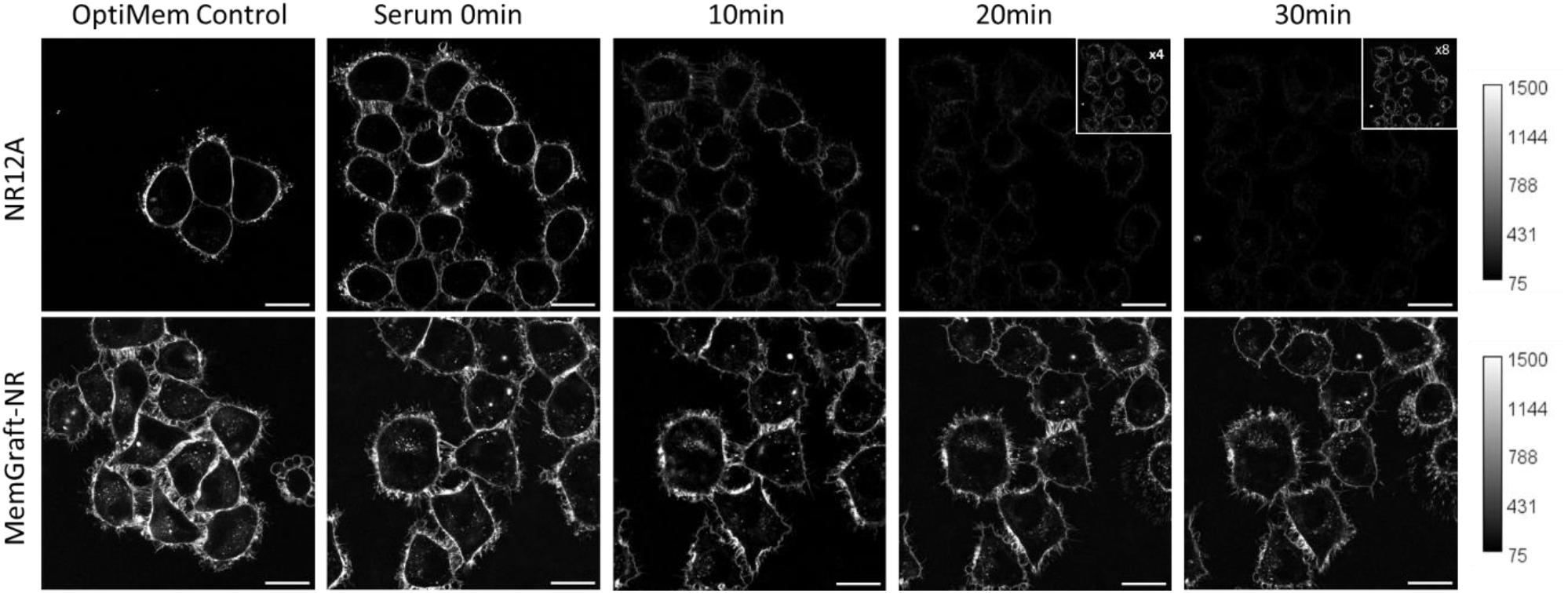
Confocal microscopy images of live KB cells stained with MemGraft-NR or reference probe NR12A, left in full serum media with images taken every 10 minutes. Excitation wavelength was 488 nm. Scale bars: 20 µm.

We also tested the resistance of MemGraft-NR to fixation and permeabilization, which are frequently used in biological experiments. To this end, the cells were stained with MemGraft-NR or a reference dye NR12A, fixed using paraformaldehyde, and then permeabilized with Tween-20 surfactant (Figure S2). MemGraft-NR resisted well to both fixation and permeabilization protocol, presenting clear PM staining. In contrast, the reference dye NR12A resisted only to fixation, but the PM labelling was compromised after permeabilization (Figure S2). Indeed, gentle fixation is not expected to impact the lipid bilayer of the PM, which allows NR12A to preserve PM labelling. However, permeabilization leads to removal of lipids and amphiphilic probes from the PM by the detergent, which explains the loss of NR12A signal. The resistance of MemGraft-NR to permeabilization is in agreement with covalent grafting of this probe at the level of membrane proteins.

### Sensing lipid surrounding of proteins by MemGraft-NR

Solvatochromic properties of MemGraft-NR were then studied in solvents of varying polarity and in the mixtures of apolar dioxane with polar methanol. Absorption and fluorescence bands showed characteristic red shifts with increase in solvent polarity (Figures S3 and S4). The positions of the band maxima were similar to those of Nile Red (Table S1), which indicates that functionalization of Nile Red with double-MemGraft scaffold did not have an impact on solvatochromism of the fluorophore itself. This was expected because all modifications were done at the aliphatic part of the fluorophore.

The solvatochromic nature of the Nile Red fluorophore allows us to explore the local environment of the probe at the grafting site. Based on our previously established methodology,^6^ the emission of the probe under the microscope was split to green (550-600 nm) and red (600-650 nm) channels, which corresponds to the two sides of the probe emission spectrum. Here, the ratio of two emission channels can reflect the polarity-dependent shifts of the probe’s emission. Since a simple ratio is non-linear, we replaced it with a function that divides the intensity of the red channel by the sum of the two channels (*I*_red_/(*I*_green_ + *I*_red_). This parameter would vary between 0 and 1, where the increase in the ratio value corresponds to an increase in polarity.

In order to analyze the molecular environment of the new MemGraft-NR probe, the ratiometric analysis was done in comparison with reference plasma membrane probe NR12A based on Nile Red. The ratiometric image of MemGraft-NR revealed PM with yellow-green pseudo-color, which corresponded to the intensity ratio of 0.55 (Figure 4). On the other hand, NR12A showed blue-green pseudo-color with the average ratio value of 0.40. Thus, the new probe experiences significantly higher local polarity, compared to NR12A. One should note that NR12A localizes in the lipid part of the outer leaflet of the PM,^6^ whereas MemGraft-NR is grafted to membrane protein of this outer leaflet. This difference in the probe localization may explain significantly higher local polarity detected by MemGraft-NR in the labelling sites of membrane proteins.

**Figure 4.**
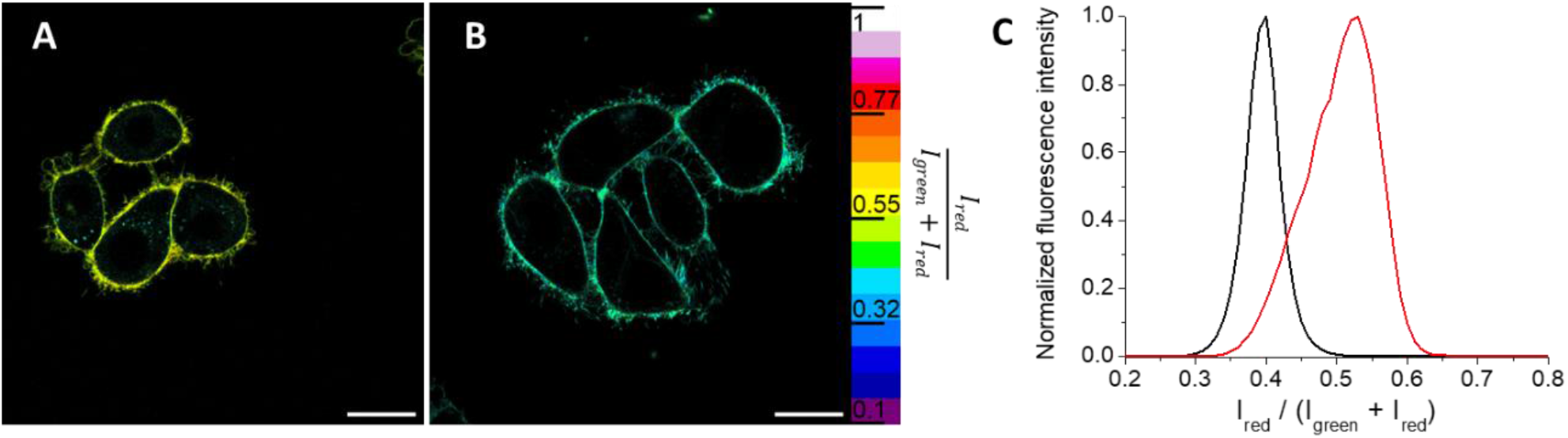
Confocal ratiometric microscopy of live KB cells stained with MemGraft-NR (A) or NR12A (B). Pseudo color reflects apparent polarity by taking the ratio of the red channel over the total intensity (red + green channels). Excitation wavelength was 488 nm. Scale bars 20 µm. (C) Average ratio distribution at the membrane of MemGraft-NR (red) and NR12A (black) stained sample.

To understand whether MemGraft-NR can still detect lipid surrounding at the grafted site of the membrane protein, we modified the PM lipid composition and compared the results with NR12A. First, we extracted cholesterol from the PM using methyl-β-cyclodextrin (MβCD) and added excess cholesterol using cholesterol-MβCD complex. We found that the removal of cholesterol leads to significant change in the pseudo-color that corresponds to the increase in the intensity ratio for both NR12A and MemGraft-NR (Figure 5). Thus, cholesterol removal decreases the lipid order, causing increase in the local polarity, in line with previous studies with NR12A and other solvatochromic membrane probes.^6^ On the other hand, the effects of cholesterol addition, which is expected to increase the local lipid order, were quite different for the two probes. Thus, NR12A showed only minor decrease in the ratio values, which were not statistically significant (Figure 5). In contrast, MemGraft-NR displayed a significant drop in the ratio on cholesterol addition, which was as strong as the effect of cholesterol extraction. These results allow us to make several key conclusions. First, MemGraft-NR can sense changes in the lipid order, suggesting that the grafted probe is exposed to lipid surrounding of the labelled protein. Second, the sensing profile of MemGraft-NR is different from that of lipid-bound NR12A when it comes to cholesterol enrichment. NR12A is poorly sensitive to the addition of cholesterol,^6^ because the PM is already saturated with cholesterol,^37^ with content values reaching 40-50 %.^38-39^ However, excess of cholesterol can significantly impact the lipid surrounding of the membrane proteins, which may be less saturated with cholesterol. Indeed, a large fraction of membrane proteins are known to avoid liquid ordered phases^40-42^ and thus present lipid surrounding with lower local cholesterol content. Then, cholesterol enrichment may impact strongly this environment, which can be reported by the probe as a strong decrease in the local polarity of the membrane protein.

**Figure 5.**
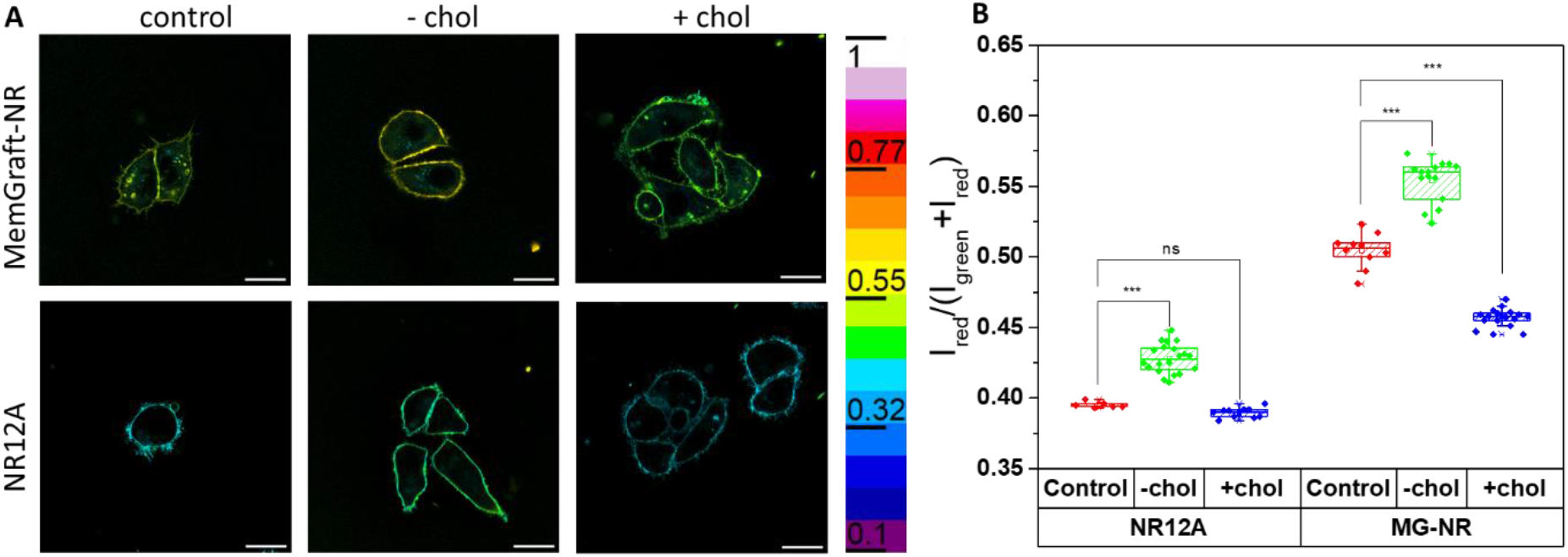
Confocal ratiometric microscopy images of live KB cells stained with MemGraft-NR or NR12A, then treated with MβCD or MβCD/chol complex (A) and average mean ratios per cell for multiple samples (B). Excitation wavelength was 488 nm. Scale bars: 20 µm. *** = p<0.001.

Second, we modified the cellular PM by exchanging native lipids with lipids of varied unsaturation level: POPC and DOPC having one and two double bonds per lipid, respectively. As the outer leaflet of PM is highly ordered due to strong content in the saturated lipids,^42^ we expect that the addition of unsaturated lipids could decrease the local lipid order. Methyl-α-cyclodextrin (MαCD) was used for the lipid exchange at the plasma membranes,^43^ owing to its capacity to bind lipids without extracting sterols, like cholesterol, in contrast to MβCD. The presence of MαCD caused a small but significant decrease in the local polarity (Figure 6), suggesting that MαCD preferentially extracts some of more “polar” lipids from the PM. Then, lipid exchange for POPC and especially for DOPC led to significant increase in the local polarity compared to control cells (Figure 6). Thus, insertion of unsaturated lipids into PM induces an expected decrease in the lipid order, which is observed as polarity increase in the protein surrounding. The effect grows with level of unsaturation from POPC and DOPC. The fact that POPC also produces a decrease in the lipid order suggests that the lipid unsaturation of the surrounding of membrane proteins is lower than that of POPC.

**Figure 6.**
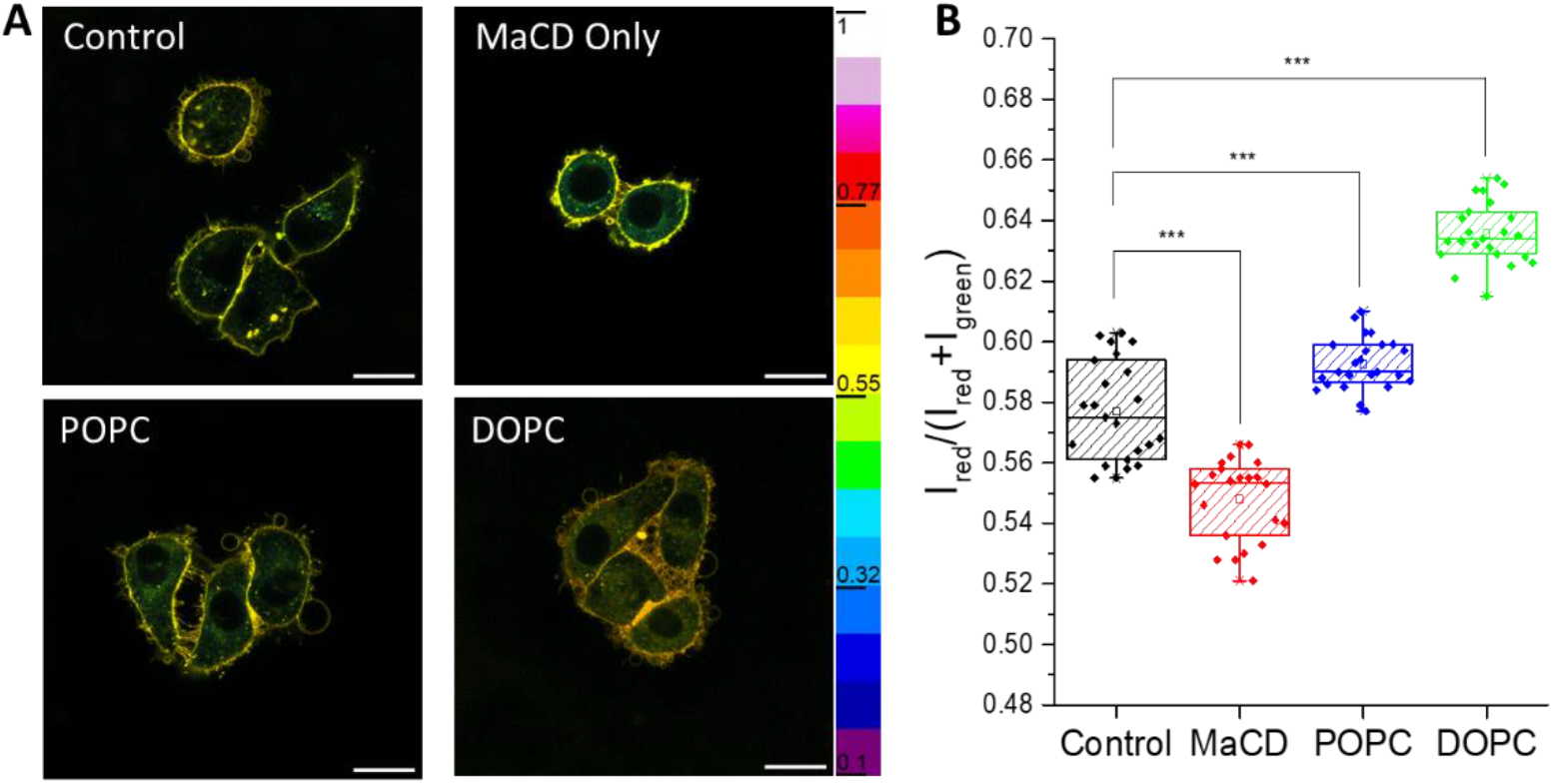
(A) Confocal ratiometric microscopy images of fixed KB cells stained with MemGraft-NR. Excitation wavelength was 488 nm. Scale bars: 20 µm. Different lipids were added using MαCD as a carrier. (B) Box plot of mean ratio values for different treatments. *** = p<0.001.

As MemGraft-NR is covalently attached to membrane proteins, it could provide unique information about lipid removal from PM during the permeabilization process. To this end, cells labelled with MemGraft-NR were fixed and then permeabilized by Tween 20. Fluorescence ratiometric images showed some changes in the emission color on permeabilization with Tween 20, while the ratio analysis revealed gradual increase in the local polarity for the first 10 minutes of the permeabilization, followed by a stabilization of the ratio (Figure 7A,B). Thus, as expected removal of lipids from protein surrounding led to the increase in the local polarity. Moreover, the process was found to take place within 10 min. Thus, MemGraft-NR enables unique observation of removal of lipids from the protein surrounding and access to the kinetics of this process, the information impossible to access with non-covalent probes. However, one should note that the changes are rather small compared to what one would expect for dye exposed to aqueous medium after lipid removal. We could speculate that during permeabilization, the lipids are replaced with the detergent, so that environment of membrane proteins remain rather apolar.

**Figure 7.**
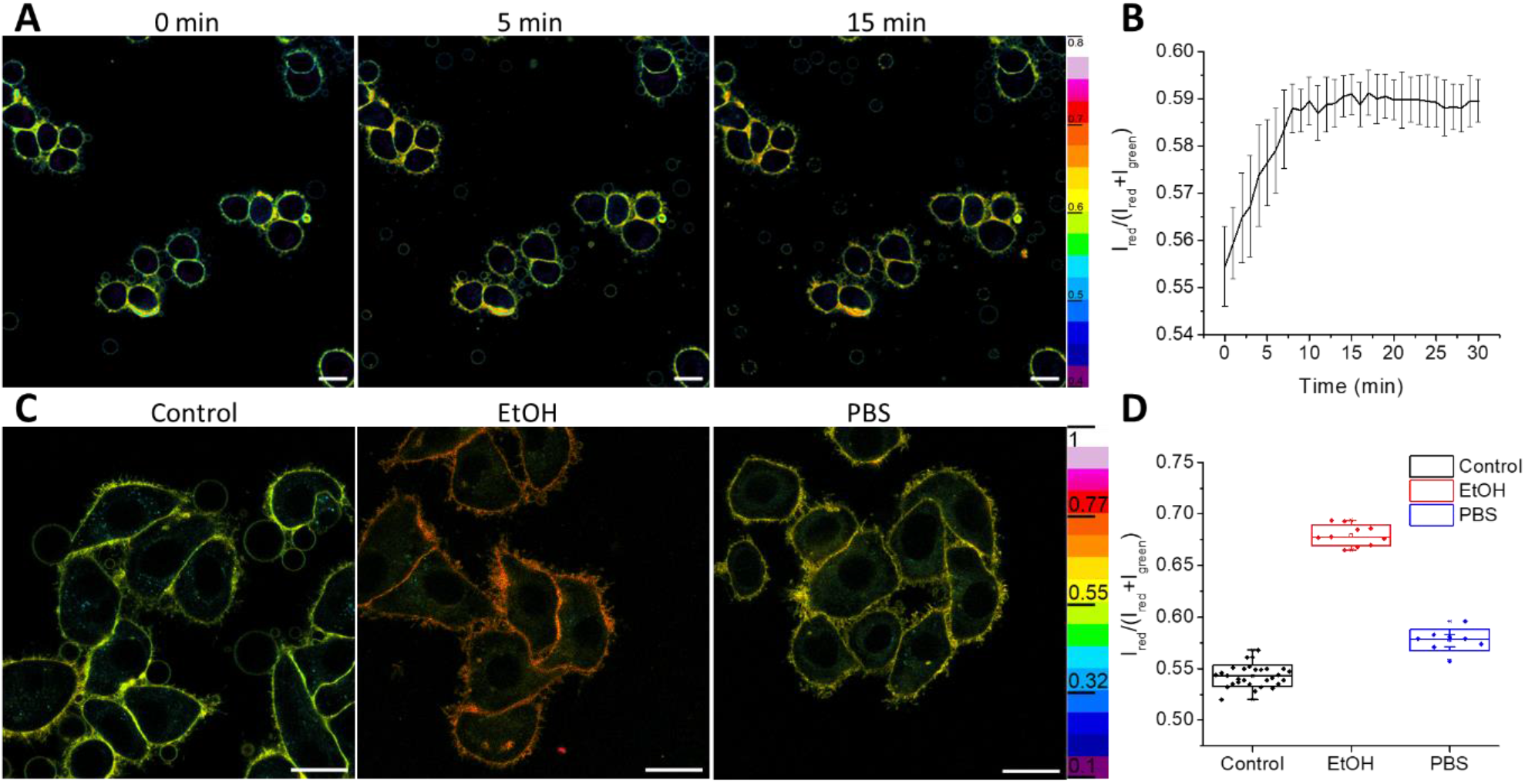
Confocal ratiometric microscopy images of fixed KB cells stained with MemGraft-NR in different conditions of permeabilization. (A) In permeabilizing solution of Tween 20 for different incubation time at RT. (B) Average ratios plotted over time. (C) Control fixed cells before (control) and after addition of 50% of ethanol (ethanol) and cell after replacing ethanol solution with PBS. (D) Average ratios plotted for the three studied conditions. Excitation wavelength was 488 nm. Scale bars (A,C): 20 µm.

We further studied effect of fixation and permeabilization with the use of organic solvents, like ethanol. After fixation with PFA, 50% of ethanol in water was added and ratiometric images were taken. Remarkably, a dramatic increase in the local polarity was observed suggesting that ethanol replaces lipids in the local protein environment (Figure 7C,D). Further displacement of the medium from ethanol to PBS led to recovery of low polarity, although the local polarity was still higher compared to freshly fixed cells (Figure 7C,D). Importantly, the final polarity values matched closely those after permeabilization with Tween 20 (Figure 7B,D). We can conclude that treatments with Tween 20 and 50% ethanol led to similar final result in terms of local polarity of membrane proteins. On the other hand, the MemGraft-NR probe also helped us to visualize the local action of ethanol, witnessing infiltration of ethanol solvent into the membrane next to proteins, something impossible to do with existing solvatochromic membrane probes.

### Monitoring endocytosis

As MemGraft-NR labels membrane proteins, it is expected to internalize inside the cells following classical endocytosis processes. We compared the internalization of MemGraft-NR to lipid probe NR12A. During incubation in reduced serum media, with NR12A we observed very slow appearance of fluorescence inside the cells (Figure 8). Only after 4 hours did we start seeing significant fluorescence in the interior of cells. With MemGraft-NR, we observed fluorescence in the cells interior already after 1h incubation (Figure 8). Quantitative analysis confirmed significantly faster accumulation of MemGraft-NR inside the cells compared to NR12A (Figure S5). Co-staining with LysoTracker Deep Red after 3h incubation of MemGraft-NR showed their good colocalization (Figure S6), which confirmed the endocytic pathway of their internalization. Thus, we can conclude that grafting to membrane proteins favors faster endocytosis compared to lipid-bound probe. Generally, lipids undergo internalization with a half-life of ∼5 min,^44^ but they are rapidly recycled back to the PM, which probably ensures long retention of NR12A on the cell surface. On the other hand, internalization of proteins is species-dependent, with varied rates of internalization and recycling.^45-47^ Therefore, some proteins get trapped in endosome/lysosomes more than others and thus tend to accumulate faster inside the cells. This could explain why MemGraft-NR presents more efficient internalization than lipid bound NR12A.

**Figure 8.**
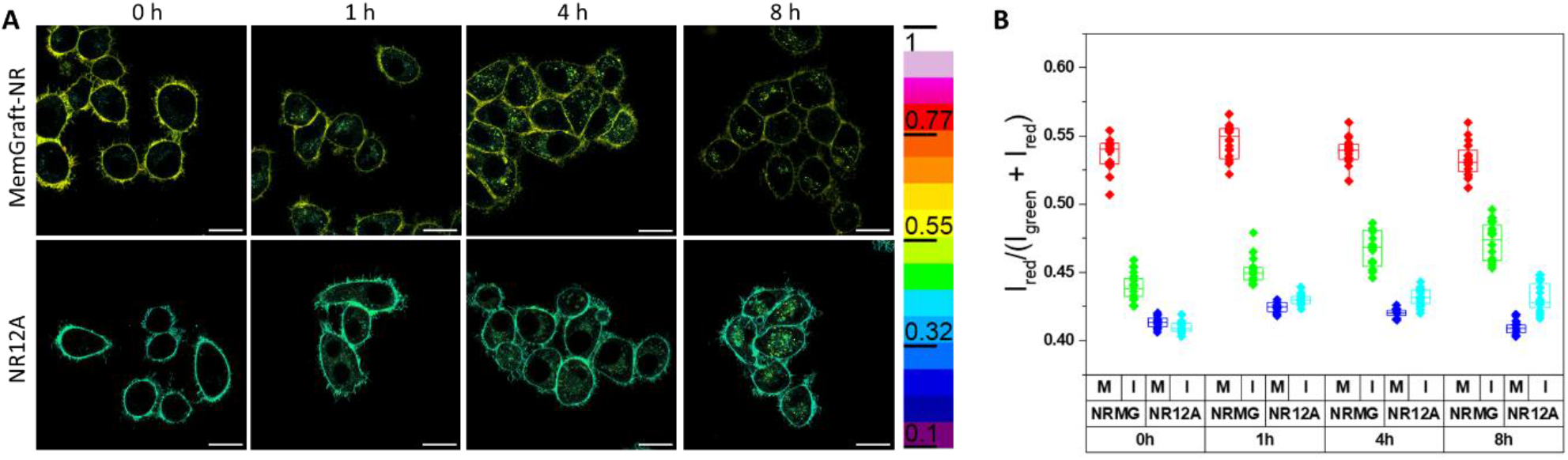
Confocal ratiometric microscopy images of live KB cells stained with MemGraft-NR or NR12A in reduced serum media, pseudo-colored to reflect the apparent polarity (A). Same sample, left in incubator between measurements. Excitation wavelength was 488 nm. Scale bars: 20 µm. (B) Box charts representing mean ratio values of membranes (M) and cell interiors (I) for MemGraft-NR (MG-NR) and NR12A and over time.

Furthermore, the ratio analysis revealed further key difference between the endocytosis of protein and lipid bound probes. Thus, in the case of NR12A, initially formed endosomes presented nearly identical local polarity as that of the PM (Figure 8B). Then, the local polarity of NR12A in endosomes slowly increased over time, whereas polarity at the PM remained virtually the same over time. In case of MemGraft-NR, the emission ratio at PM also remained constant over time (Figure 8B). One should note that even longer incubation times over 24h did not change the ratio at the PM of MemGraft-NR (Figure S1), which suggests that the cells maintain constant lipid organization at the interface with membrane proteins. In contrast, the newly formed endosomes showed a significantly lower polarity compared to the PM. Over time they underwent continuous increase in polarity for both MemGraft-NR and NR12A (Figure 8), in line with previous observation for analogous probe NR12S.^48^ This increase in the polarity is connected with endosomal maturation, which is generally associated with the removal of cholesterol and sphingomyelin from the endosome,^49-50^ which leads to the decrease in the local lipid order of the endosomal membranes. Much more surprising is the observed much lower polarity in the newly formed endosomes (Figure 8), which are expected to present the same protein-lipid composition to that of the PM. It is possible that the high negative curvature of endosomes inner leaflet may immerse the labelling site membrane proteins deeper into the lipid bilayer and/or bring proteins closer, thus decreasing the local polarity. The absence of this effect with NR12A indicates that endosome formation alters the organization at the protein– lipid interface rather than the organization of the lipid membrane itself. Thus, sensing lipid surroundings of proteins reveals significant changes at the protein-lipid interface of biomembranes, which were not visible before with classical non-covalent probes.

## Conclusion

Organization of lipids in cell plasma membranes have been traditionally studied using lipid-like probes sensitive to the environment. In this work, we propose an original probe design that enables sensing lipid organization in the immediate surrounding of proteins, i.e. at the protein-lipid interface. This was achieved by applying a concept of lipid-directed protein labeling, recently introduced by us, with solvatochromic dye Nile Red, which has been proven to be a powerful tool to study lipid membrane organization. To this end, we designed a new MemGraft scaffold combining together (i) a low affinity membrane anchor, built of a charged sulfonate group and a short alkyl chain (butyl), (ii) amino group coupled to a fluorescent dye and (iii) carboxylate in form of NHS active ester. While the membrane anchor is expected to ensure reversible docking of the probe to the lipid membrane, the NHS ester is responsible for grafting the environment-sensitive probe to the amino groups of proteins. However, the Nile Red derivative bearing this scaffold (MemGraft-NR v1) was insufficient to ensure robust PM labelling. Therefore, we synthesized Nile Red bearing a sequence of two MemGraft scaffolds. The new probe, MemGraft-NR v2 (MemGraft-NR) showed excellent labelling of plasma membranes, similar to that of reference membrane probes. Unlike classical lipid-like probe based on Nile Red (NR12A) it showed high resistance to the presence of serum and fixation with permeabilization, owing to the covalent nature of the labelling. Taking advantage of the ability of Nile Red to sense lipid organization in biomembranes, MemGraft-NR was studied by two-color confocal imaging in live and fixed cells. We found that the local environment at the protein lipid interface of MemGraft-NR is more polar than that of lipid probe NR12A. At the same time, MemGraft-NR remained sensitive to variation of cholesterol content in PM of live cells, which confirmed the probe being grafted to membrane proteins detects their local lipid surrounding. Surprisingly, its sensitivity to cholesterol variation was even higher compared to NR12A, suggesting that protein-lipid interface is an important region in the PM, highly sensitive to variations in lipid composition. The new probe allowed membrane sensing that were not possible before with classical solvatochromic PM probes: (1) in situ sensing changes in the lipid exchange, which revived strong polarity increase with PM enrichment with unsaturated lipid DOPC; (2) dynamic monitoring of permeabilization processes, revealing that lipid removal associated with local polarity rise takes place in less than 10 min; (3) dramatic changes in the local protein surrounding (polarity jump) in the presence of 50% ethanol, which also further leads to cell permeabilization. Finally, we found that the MemGraft-NR internalizes by endocytosis a bit faster than corresponding lipid probe, highlighting that peculiar difference in the internalization of protein and lipid bound molecules. Moreover, a dramatic decrease in the local polarity was revealed by MemGraft-NR at the initial steps of endocytosis, in contrast to lipid-based probe, suggesting that the negative curvature of endosomes provide dramatic effect on organization of the protein-lipid interface. We can speculate that the labelling site of proteins at the interface with lipids gets deeper imbedded into lipid membrane at the inner leaflet of endosomes due to negative curvature, thus triggering decrease in the local polarity. Overall, we propose a new tool to study biological membrane that drastically differ from those reported before, because it senses biophysical properties at the protein-lipid interface. The approach to sense lipid surrounding of proteins appears highly promising because its properties, sensitivity to external stimuli and endocytosis are different from those observed in lipid part of plasma membranes.

## Materials and methods

All the reagents were purchased from Sigma-Aldrich, Alfa Aesar, Fisher Scientific or TCI and used without any purification. MilliQ-water (Millipore) was used in all experiments. NMR spectra were recorded on a Bruker Avance III 400 MHz spectrometer and a Bruker Avance III 500 MHz spectrometer. Mass spectra were obtained using an Agilent Q-TOF 6520 mass spectrometer with electrospray ionization. NR-COOH was synthesized as previously reported.^6^

### Synthesis of probes

#### Compound 1: 3-(butylammonio)propane-1-sulfonate

1,3-propanesultone (1 eq. 16 mmol, 2 g) was dissolved in dry MeOH under Argon. n-Butylamine (2 eq. 32 mmol, 3.2 mL) was added dropwise to the solution. The reaction was left to stir overnight. The solvent was evaporated in vacuo and acetone was added to wash the solid compound. The solid was filtered out and dried to afford compound **1** (2.66g, 85%). ^1^H NMR corresponded to those reported elsewhere.^6 1^H NMR (400 MHz, D_2_O) δ 3.23 – 3.09 (m, 2H), 3.07 – 2.85 (m, 4H), 2.17 – 1.98 (m, 2H), 1.70 – 1.53 (m, 2H), 1.45 – 1.28 (m, 2H), 0.92 (t, J = 7.4 Hz, 3H) ppm.

#### Compound 2: Sodium 3-(2-((((9H-fluoren-9-yl)methoxy)carbonyl)amino)-4-(tert-butoxy)-N-butyl-4-oxobutanamido)propane-1-sulfonate

Fmoc-Asp(OtBu)-OH (1 eq., 2.33 mmol, 0.96 g) and **1** (1.1 eq., 2.56 mmol, 0.5 g) were dissolved in 12 mL of dry DMF under Argon. DIPEA (3.5 eq., 8.16 mmol, 1.42 mL) was added to the stirring solution. Then PyBOP (1.1 eq., 2.56 mmol, 1.33 g) was added. The solution was stirred overnight at room temperature. DMF was evaporated in vacuo and the crude was purified using column chromatography (89.5:10:0.5 v/v/v DCM:MeOH:TEA). The product was taken up in ethyl acetate and washed 3x with brine. Ethyl acetate was evaporated in vacuo to yield product **2** as an off-white solid (733 mg, 51%). ^1^H NMR (400 MHz, MeOD): δ 7.82 – 7.71 (m, 2H), 7.71 – 7.60 (m, 2H), 7.41 –7.34 (m, 2H), 7.34 – 7.26 (m, 2H), 4.95 – 4.88 (m, 1H), 4.50 – 4.27 (m, 2H), 4.20 (q, J = 6.5 Hz, 1H), 3.68 – 3.34 (m, 4H), 2.89 – 2.69 (m, 3H), 2.48 (dt, J = 15.8, 6.5 Hz, 1H), 2.20 – 1.95 (m, 2H), 1.72 –1.48 (m, 2H), 1.42 (s, 9H), 1.38 – 1.25 (m, 2H), 0.97 – 0.87 (m, 3H). HRMS (ESI+), m/z: [M+2H]^+^ calcd for C_30_H_41_N_2_O_8_S^+^, 589.2578; found, 589.2578

#### Compound 3: Sodium 3-(2-amino-4-(tert-butoxy)-N-butyl-4-oxobutanamido)propane-1-sulfonate

**2** (1 eq., 0.4 mmol, 0.36g) was dissolved in dry DMF (8 mL) then piperidine (2 mL) was added and the mixture left to stir at room temperature for 30 min. Solvents were evaporated in vacuo and the solid washed using 50:50 v/v MeOH:H_2_O. The filtrate was recovered and solvents removed to afford the product **3** as a white solid (155 mg, 68%). ^1^H NMR (400 MHz, MeOD): δ 4.12 (m, 1H), 3.77 – 3.34 (m, 4H), 2.91 – 2.75 (m, 2H), 2.65 (m, 1H), 2.52 – 2.35 (m, 1H), 2.22 – 1.96 (m, 2H), 1.74 – 1.47 (m, 2H), 1.45 (m, 9H), 1.40 – 1.26 (m, 2H), 0.96 (m, 3H). HRMS (ESI+), m/z: [M+2H]^+^ calcd for C_15_H_31_N_2_O_6_S^+^, 367.1897; found, 367.1911

#### Compound 4a: NileRed-Asp(C4)-OtBu, 3-(4-(tert-butoxy)-N-butyl-2-(4-(ethyl(5-oxo-5H-benzo[a]phenoxazin-9-yl)amino)butanamido)-4-oxobutanamido)propane-1-sulfonate

NR-COOH (1 eq., 186 µmol, 70 mg) and universal anchor **3** (1.1 eq., 205 µmol, 80 mg) were dissolved in anhydrous DMF (10 mL), then PyBOP (1.1 eq., 205 µmol, 106 mg) and DIPEA (5 eq., 930 µmol, 150 µL) were added. The reaction mixture was stirred under argon for 18 hours at room temperature. The solvent was evaporated in vacuo, the residue was dissolved in EtOAC and washed with 1M HCl (3x 50 mL), then brine (50 mL) and dried over Na_2_SO_4_ and filtered, then the solvent was removed under vacuum. The crude was purified by reverse-phase flash chromatography using a gradient of ACN in H_2_O to yield **4a** as a dark purple solid (70 mg, 50%). ^1^H NMR (400 MHz, MeOD) δ 8.81 (dd, J = 13.2, 8.0 Hz, 1H), 8.64 (d, J = 7.8 Hz, 1H), 8.30 (t, J = 8.6 Hz, 1H), 8.21 (d, J = 7.8 Hz, 1H), 7.86 – 7.70 (m, 2H), 7.01 (d, J = 52.2 Hz, 1H), 6.83 (s, 1H), 5.23 (q, J = 6.6 Hz, 1H), 3.79 – 3.43 (m, 3H), 2.98 (td, J = 6.7, 3.6 Hz, 1H), 2.77 (ddt, J = 11.0, 7.5, 3.7 Hz, 2H), 2.57 – 2.44 (m, 2H), 2.42 – 2.31 (m, 2H), 2.04 – 1.99 (m, 3H), 1.73 – 1.66 (m, 2H), 1.43 (s, 9H), 1.36 – 1.21 (m, 8H), 0.99 (q, J = 7.2 Hz, 3H), 0.90 (t, J = 8.2 Hz, 3H).

#### Compound 4: NileRed-Asp(C4)-OH, 3-(N-butyl-3-carboxy-2-(4-(ethyl(5-oxo-5H-benzo[a]phenoxazin-9-yl)amino)butanamido)propanamido)propane-1-sulfonate

**4a** NR-Asp(C4)-OtBu (1 eq., 70mg) was dissolved in dry DCM (5 mL) then TFA (1 mL) was added. The mixture instantly turned blue and was left to stir at room temperature for 1 hour. The solvents were evaporated under reduced pressure. The crude product was used in the next step without further purification (quantitative yield assumed).

#### Compound 5: MemGraft-NR v1 / NileRed-Asp(C4)-Osu 3-(N-butyl-4-((2,5-dioxopyrrolidin-1-yl)oxy)-2-(4-(ethyl(5-oxo-5H-benzo[a]phenoxazin-9-yl)amino)butanamido)-4-oxobutanamido)propane-1-sulfonate

NR-Asp(C4)-OH (1 eq., 46 µmol, 32 mg) and TSTU (1.5 eq., 69 µmol, 21 mg) were dissolved in dry DCM (4 mL) then, DIPEA (1.5 eq., 69 µmol, 12 µL) was added and the mixture stirred at room temperature for 3 hours. The solvent was evaporated and the crude rapidly purified by reverse--phase flash chromatography to yield the desired product as a purple solid (5 mg, 14%). LC-MS (ESI+) m/z: [M+H+NH4]^+^ calcd for C_37_H_47_N_6_O_11_S^+^ 783.30, found 783.34

#### Compound 6a: NileRed-Asp(C4)2-OtBu

**4** NR-Asp-OH (1 eq., 94 µmol, 65 mg), **3** (1.1 eq., 103 µmol, 40 mg) and PyBOP (2 eq., 186 µmol, 96 mg) were dissolved in dry DMF (2 mL), then DIPEA (10 eq., 940 µmol, 160 µL) was added and the mixture left to stir at room temperature for 18 hours. The solvents were removed in vacuo and the resulting crude purified by reverse-phase flash chromatography to yield **6a** NR-Asp2-OtBu as a purple solid (20 mg, 21%). ^1^H NMR (500 MHz, MeOD) δ 8.59 (dd, J = 14.2, 8.1 Hz, 1H), 8.21 – 8.13 (m, 1H), 7.80 – 7.57 (m, 3H), 6.84 (t, J = 10.1 Hz, 1H), 6.59 (s, 1H), 6.30 (d, J = 16.0 Hz, 1H), 4.55 (s, 2H), 3.60 – 3.34 (m, 8H), 2.84 – 2.71 (m, 4H), 2.44 – 2.31 (m, 4H), 2.25 – 2.09 (m, 2H), 2.09 – 1.90 (m, 8H), 1.49 (dt, J = 15.1, 7.5 Hz, 4H), 1.42 (d, J = 15.2 Hz, 9H), 1.37 – 1.15 (m, 8H), 1.00 (q, J = 7.0 Hz, 3H), 0.89 (td, J = 7.6, 5.0 Hz, 6H). LC-MS (ESI-) m/z: [M]^2-^ calcd for C_48_ H_66_ N_6_ O_14_ S_2_ ^2-^ 507.20, found 507.20.

#### Compound 6: NileRed-Asp(C4)2-OH

**6a** NR-Asp2-OtBu (1 eq., 17mg) was dissolved in dry DCM (5 mL) then TFA (1 mL) was added. The mixture instantly turned blue and was left to stir at room temperature for 1 hour. The solvents were evaporated under reduced pressure. The crude product was purified by reverse-phase flash chromatography to yield the final compound (6 mg, 37%). ^1^H NMR (500 MHz, MeOD) δ 8.63 – 8.52 (m, 1H), 8.18 (s, 1H), 7.77 (d, J = 7.6 Hz, 1H), 7.67 (d, J = 7.1 Hz, 1H), 7.64 – 7.52 (m, 1H), 6.90 – 6.79 (m, 1H), 6.59 (d, J = 21.2 Hz, 1H), 6.35 (d, J = 12.8 Hz, 1H), 5.45 – 5.38 (m, 1H), 5.35 – 5.25 (m, 1H), 3.48 (d, J = 33.2 Hz, 8H), 3.28 – 3.22 (m, 2H), 3.07 – 2.71 (m, 2H), 2.63 (t, J = 12.9 Hz, 2H), 2.45 (d, J = 67.1 Hz, 2H), 2.06 – 1.90 (m, 8H), 1.66 (d, J = 8.1 Hz, 2H), 1.48 (d, J = 8.2 Hz, 3H), 1.24 (t, J = 7.5 Hz, 3H), 1.04 – 0.77 (m, 6H). LC-MS (ESI-), m/z: [M-H]-calcd. for C44H59N6O14S2-959.3536, found 959.3524.

#### Compound 7: NileRed MemGraft v2 / MGNR / NileRed-Asp(C4)2-OSu

NR-Asp2-OH (1 eq., 3 mg, 2.9 µmol) was dissolved in a 1:1 mixture of dry ACN and dry DMF (1 mL) then TSTU (2 eq., 6 µmol, 2 mg) and DIPEA (20 eq., 60 µmol, 10 µL) were added and the mixture left to stir at room temperature for 4 hours. Solvents were evaporated and the product used as it is (quant. yield assumed). LC-MS (ESI-), m/z: [M-H]-calcd for C48H62N7O16S2-1056.3700, found 1056.3695.

### Cell experiments

KB (ATCC CRL-3596) cells were grown in Eagle’s Minimum essential medium (EMEM, Gibco Invitrogen), supplemented with 10% fetal bovine serum (FBS, Lonza), 2 mM L-glutamine (Gibco-Invitrogen), 1% non-essential amino acid solution (Gibco-Invitrogen) and sodium pyruvate 1 mmol/L at 37 °C in a humidified 5% CO2 atmosphere. Cells were seeded onto a chambered coverglass (IBIDI) at a density of 5 × 10^4^ cells/well, or in 8 chamber coverglass (Lab-Tek) at 2.5 × 10^4^ cells/well, 24 h before the microscopy measurement.

For microscopy imaging, the attached live cells in IBIDI dishes were washed once with warm Hanks’ balanced salt solution (HBSS, Gibco-Invitrogen); after that, 1 mL of a corresponding MemGraft dye solution at 500 nM in HBSS was added and the cells were incubated for 5 min at room temperature. Then, the attached cells in IBIDI dishes were washed twice with HBSS before imaging in OptiMEM (Gibco-Invitrogen). Cells stained with NR12A were washed once with warm HBSS and imaged in OptiMEM with 50 nM NR12A dye.

For fixation and permeabilization experiments, the dye solution was removed from the cells and the attached cells were washed twice with HBSS and incubated for 15 min with 4% (w/v) formaldehyde solution in phosphate-buffered saline (PBS) at room temperature. The fixative solution was removed and the cells were washed by pipetting PBS against the side of the dish 3 times. In the case of NR12A, it was added after washing away the fixative solution. Then, the cells were incubated with a permeabilizing solution (0.1 % Tween 20 in PBS) under the microscope and time-lapsed.

Cholesterol extractions and additions were performed using a solution of methyl-beta-cyclodextrin (MβCD) at 5 mM in OptiMEM, or a complex of MβCD with cholesterol 2:1 at 5 mM and 2.5 mM respectively, and incubating for 1h at 37°C. The MβCD solution was then washed away with HBSS. Cells stained with MGNR were stained before MβCD treatment, and cells stained with NR12A were stained after MβCD treatment as multiple washing steps will remove NR12A.

Lipid replacements were performed using methyl-alpha-cyclodextrin (MαCD, Cyclodextrin Shop) at a concentration of 40 mM in PBS. Lipid-MαCD complexes were prepared by first drying the lipids (AvantiLipids), slowly hydrating them with the solution of 40 mM MαCD in PBS and then vortexing for a final lipid concentration of 1.5 mM. Cells stained with MGNR were fixed and incubated with the corresponding lipid-MαCD solution, or MαCD only, during 1 hour at 37°C. The lipid solution was then removed and cells washed twice with PBS and imaged in PBS.

### Fluorescence Microscopy

Confocal imaging of cells was performed on a Leica TCS SP8 confocal microscope with a HCX PL APO 63×/ 1.40 OIL CS2 objective and two 12-bit photomultipliers. The excitation light was provided by lasers of 488 nm, and 638 nm. All the parameters at each channel were left constant; the illumination power was adjusted to achieve a good signal for each probe. All the images were processed using ImageJ software.

### Ratiometric imaging

For NileRed ratiometric imaging, the samples were illuminated with 488 nm laser, the green channel was defined at 550-600 nm and the red channel at 600-650 nm. RatioloJ plugin for ImageJ (developed by Romain Vauchelles), distributed by PIQ-QuESt platform (https://piq.unistra.fr/) from university of Strasbourg, was used to analyze the images using the following intensity ratio: Intensity in red channel / (Intensity in green channel + Intensity in red channel).

### Statistical analysis

Statistical tests were performed Shapiro-Wilk tests were used on all data sets to confirm normal distribution. Statistical analyses were performed using one-way ANOVA with significance set at α = 0.001 for Tukey means comparison.

## Supporting information

Supplemental information for publication.

## Acknowledgements

This work was supported by Agence Nationale de la Recherche (ANR) EV-Graft ANR-24-CE44-4520-01. TLCM is supported by a Contrat Doctoral Spécifique Normalien (CDSN) grant from the French Ministry of Higher Education and Research (MESR), distributed by ENS-PSL. We acknowledge PIQ platform of France-Bioimaging (FBI) network for providing access to fluorescence imaging facilities and PACSI platform − for NMR and mass spectrometry.

## References

(1) Stillwell, W. An Introduction to Biological Membranes: Composition, Structure and Function; Elsevier Science, 2016.

(2) Sezgin, E.; Levental, I.; Mayor, S.; Eggeling, C. The Mystery of Membrane Organization: Composition, Regulation and Roles of Lipid Rafts. Nat Rev Mol Cell Biol 2017, 18, 361–374.

(3) Lingwood, D.; Simons, K. Lipid Rafts as a Membrane-Organizing Principle. Science 2010, 327, 46–50.

(4) Klymchenko, A. S. Fluorescent Probes for Lipid Membranes: From the Cell Surface to Organelles. Acc Chem Res 2023, 56, 1–12.

(5) Liu, C.; Gao, X.; Yuan, J.; Zhang, R. Advances in the Development of Fluorescence Probes for Cell Plasma Membrane Imaging. TrAC Trends in Analytical Chemistry 2020, 133, 116092.

(6) Danylchuk, D. I.; Moon, S.; Xu, K.; Klymchenko, A. S. Switchable Solvatochromic Probes for Live-Cell Super-Resolution Imaging of Plasma Membrane Organization. Angewandte Chemie International Edition 2019, 58, 14920–14924.

(7) Kucherak, O. A.; Oncul, S.; Darwich, Z.; Yushchenko, D. A.; Arntz, Y.; Didier, P.; Mély, Y.; Klymchenko, A. S. Switchable Nile Red-Based Probe for Cholesterol and Lipid Order at the Outer Leaflet of Biomembranes. Journal of the American Chemical Society 2010, 132, 4907–4916.

(8) Collot, M.; Ashokkumar, P.; Anton, H.; Boutant, E.; Faklaris, O.; Galli, T.; Mély, Y.; Danglot, L.; Klymchenko, A. S. Membright: A Family of Fluorescent Membrane Probes for Advanced Cellular Imaging and Neuroscience. Cell Chemical Biology 2019, 26, 600-614.e607.

(9) Dal Molin, M.; Verolet, Q.; Colom, A.; Letrun, R.; Derivery, E.; Gonzalez-Gaitan, M.; Vauthey, E.; Roux, A. l.; Sakai, N.; Matile, S. Fluorescent Flippers for Mechanosensitive Membrane Probes. Journal of the American Chemical Society 2015, 137, 568–571.

(10) López-Duarte, I.; Vu, T. T.; Izquierdo, M. A.; Bull, J. A.; Kuimova, M. K. A Molecular Rotor for Measuring Viscosity in Plasma Membranes of Live Cells. Chemical Communications 2014, 50, 5282–5284.

(11) Shimomura, T.; Seino, R.; Umezaki, K.; Shimoda, A.; Ezoe, T.; Ishiyama, M.; Akiyoshi, K. New Lipophilic Fluorescent Dyes for Labeling Extracellular Vesicles: Characterization and Monitoring of Cellular Uptake. Bioconjugate Chemistry 2021, 32, 680–684.

(12) Ling, J.; Liu, Y.; Dumoulin, A.; Sheng, D.; Fu, Y.; Liu, S.; Ding, L.; Huang, L.; Xi, P.; Tang, H.; Stoeckli, E. T.; Chen, Z. A Gentle Palette of Plasma Membrane Dyes. Proceedings of the National Academy of Sciences 2025, 122, e2504879122.

(13) Wang, W.; Zhang, Y.; Zhao, H.; Zhuang, X.; Wang, H.; He, K.; Xu, W.; Kang, Y.; Chen, S.; Zeng, S.; Qian, L. Real-Time Imaging of Cell-Surface Proteins with Antibody-Based Fluorogenic Probes. Chemical Science 2021, 12, 13477–13482.

(14) Balčiūnaitė-Murzienė, G.; Dzikaras, M. Wheat Germ Agglutinin—from Toxicity to Biomedical Applications. Applied Sciences 2021, 11, 884.

(15) Drew, D. E.; von Heijne, G.; Nordlund, P.; de Gier, J. W. Green Fluorescent Protein as an Indicator to Monitor Membrane Protein Overexpression in Escherichia Coli. FEBS Lett 2001, 507, 220–224.

(16) Roßmann, K.; Pabst, U.; Baciu, B. C.; Sun, S.; Huhn, C.; Olesen, C. H.; Kowald, M.; Tapp, E.; Bieck, M.; Birke, R.; Shields, B. C.; Jeong, P.; Hong, J.; Tadross, M. R.; Levitz, J.; Lehmann, M.; Lipstein, N.; Broichhagen, J. A General One-Step Protocol to Generate Impermeable Fluorescent Halotag Substrates for in Situ Live Cell Application and Super-Resolution Imaging. Nature Communications 2026, 17, 426.

(17) Prifti, E.; Reymond, L.; Umebayashi, M.; Hovius, R.; Riezman, H.; Johnsson, K. A Fluorogenic Probe for Snap-Tagged Plasma Membrane Proteins Based on the Solvatochromic Molecule Nile Red. ACS Chemical Biology 2014, 9, 606–612.

(18) Pelletier, R. m.; Danylchuk, D. I.; Benaissa, H.; Broch, F.; Vauchelles, R.; Gautier, A.; Klymchenko, A. S. Genetic Targeting of Solvatochromic Dyes for Probing Nanoscale Environments of Proteins in Organelles. Analytical Chemistry 2023, 95, 8512–8521.

(19) Straková, K. n.; López-Andarias, J.; Jiménez-Rojo, N.; Chambers, J. E.; Marciniak, S. J.; Riezman, H.; Sakai, N.; Matile, S. Haloflippers: A General Tool for the Fluorescence Imaging of Precisely Localized Membrane Tension Changes in Living Cells. ACS Central Science 2020, 6, 1376–1385.

(20) Swiecicki, J. M.; Santana, J. T.; Imperiali, B. A Strategic Approach for Fluorescence Imaging of Membrane Proteins in a Native-Like Environment. Cell Chem Biol 2020, 27, 245-251.e243.

(21) Kang, K.; Joo, S.; Choi, J. Y.; Geum, S.; Hong, S.-P.; Lee, S.-Y.; Kim, Y. H.; Kim, S.-M.; Yoon, M.-H.; Nam, Y.; Lee, K.-B.; Lee, H.-Y.; Choi, I. S. Tissue-Based Metabolic Labeling of Polysialic Acids in Living Primary Hippocampal Neurons. Proceedings of the National Academy of Sciences 2015, 112, E241–E248.

(22) Dumont, A.; Malleron, A.; Awwad, M.; Dukan, S.; Vauzeilles, B. Click-Mediated Labeling of Bacterial Membranes through Metabolic Modification of the Lipopolysaccharide Inner Core. Angewandte Chemie International Edition 2012, 51, 3143–3146.

(23) Fujishima, S.-h.; Yasui, R.; Miki, T.; Ojida, A.; Hamachi, I. Ligand-Directed Acyl Imidazole Chemistry for Labeling of Membrane-Bound Proteins on Live Cells. Journal of the American Chemical Society 2012, 134, 3961–3964.

(24) Nonaka, H.; Sakamoto, S.; Shiraiwa, K.; Ishikawa, M.; Tamura, T.; Okuno, K.; Kondo, T.; Kiyonaka, S.; Susaki, E. A.; Shimizu, C.; Ueda, H. R.; Kakegawa, W.; Arai, I.; Yuzaki, M.; Hamachi, I. Bioorthogonal Chemical Labeling of Endogenous Neurotransmitter Receptors in Living Mouse Brains. Proceedings of the National Academy of Sciences 2024, 121, e2313887121.

(25) Aknine, N.; Pelletier, R.; Klymchenko, A. S. Lipid-Directed Covalent Labeling of Plasma Membranes for Long-Term Imaging, Barcoding and Manipulation of Cells. JACS Au 2025, 5, 922–936.

(26) Levental, I.; Lyman, E. Regulation of Membrane Protein Structure and Function by Their Lipid Nano-Environment. Nat Rev Mol Cell Biol 2023, 24, 107–122.

(27) Klymchenko, A. S. Solvatochromic and Fluorogenic Dyes as Environment-Sensitive Probes: Design and Biological Applications. Acc Chem Res 2017, 50, 366–375.

(28) Reichardt, C. Solvatochromic Dyes as Solvent Polarity Indicators. Chemical Reviews 2002, 94, 2319–2358.

(29) Paez-Perez, M.; Kuimova, M. K. Molecular Rotors: Fluorescent Sensors for Microviscosity and Conformation of Biomolecules. Angewandte Chemie International Edition 2024, 63, e202311233.

(30) Haidekker, M. A.; Theodorakis, E. A. Molecular Rotors—Fluorescent Biosensors for Viscosity and Flow. Organic & Biomolecular Chemistry 2007, 5, 1669–1678.

(31) Colom, A.; Derivery, E.; Soleimanpour, S.; Tomba, C.; Molin, M. D.; Sakai, N.; González-Gaitán, M.; Matile, S.; Roux, A. A Fluorescent Membrane Tension Probe. Nat Chem 2018, 10, 1118–1125.

(32) Owen, D. M.; Rentero, C.; Magenau, A.; Abu-Siniyeh, A.; Gaus, K. Quantitative Imaging of Membrane Lipid Order in Cells and Organisms. Nat Protoc 2011, 7, 24–35.

(33) Klymchenko, A. S.; Kreder, R. Fluorescent Probes for Lipid Rafts: From Model Membranes to Living Cells. Chem Biol 2014, 21, 97–113.

(34) Dietrich, C.; Bagatolli, L. A.; Volovyk, Z. N.; Thompson, N. L.; Levi, M.; Jacobson, K.; Gratton, E. Lipid Rafts Reconstituted in Model Membranes. Biophysical Journal 2001, 80, 1417–1428.

(35) Jin, L.; Millard, A. C.; Wuskell, J. P.; Dong, X.; Wu, D.; Clark, H. A.; Loew, L. M. Characterization and Application of a New Optical Probe for Membrane Lipid Domains. Biophysical Journal 2006, 90, 2563–2575.

(36) Niko, Y.; Didier, P.; Mely, Y.; Konishi, G.-i.; Klymchenko, A. S. Bright and Photostable Push-Pull Pyrene Dye Visualizes Lipid Order Variation between Plasma and Intracellular Membranes. Scientific Reports 2016, 6, 18870.

(37) Das, A.; Brown, M. S.; Anderson, D. D.; Goldstein, J. L.; Radhakrishnan, A. Three Pools of Plasma Membrane Cholesterol and Their Relation to Cholesterol Homeostasis. Elife 2014, 3.

(38) van Meer, G.; Voelker, D. R.; Feigenson, G. W. Membrane Lipids: Where They Are and How They Behave. Nat Rev Mol Cell Biol 2008, 9, 112–124.

(39) Lange, Y.; Swaisgood, M. H.; Ramos, B. V.; Steck, T. L. Plasma Membranes Contain Half the Phospholipid and 90% of the Cholesterol and Sphingomyelin in Cultured Human Fibroblasts. J Biol Chem 1989, 264, 3786–3793.

(40) Schlebach, J. P.; Barrett, P. J.; Day, C. A.; Kim, J. H.; Kenworthy, A. K.; Sanders, C. R. Topologically Diverse Human Membrane Proteins Partition to Liquid-Disordered Domains in Phase-Separated Lipid Vesicles. Biochemistry 2016, 55, 985–988.

(41) Baumgart, T.; Hammond, A. T.; Sengupta, P.; Hess, S. T.; Holowka, D. A.; Baird, B. A.; Webb, W. W. Large-Scale Fluid/Fluid Phase Separation of Proteins and Lipids in Giant Plasma Membrane Vesicles. Proceedings of the National Academy of Sciences 2007, 104, 3165–3170.

(42) Lorent, J. H.; Levental, K. R.; Ganesan, L.; Rivera-Longsworth, G.; Sezgin, E.; Doktorova, M.; Lyman, E.; Levental, I. Plasma Membranes Are Asymmetric in Lipid Unsaturation, Packing and Protein Shape. Nat Chem Biol 2020, 16, 644–652.

(43) Li, G.; Kim, J.; Huang, Z.; St Clair, J. R.; Brown, D. A.; London, E. Efficient Replacement of Plasma Membrane Outer Leaflet Phospholipids and Sphingolipids in Cells with Exogenous Lipids. Proc Natl Acad Sci U S A 2016, 113, 14025–14030.

(44) Tanaka, M.; Kikuchi, T.; Uno, H.; Okita, K.; Kitanishi-Yumura, T.; Yumura, S. Turnover and Flow of the Cell Membrane for Cell Migration. Scientific Reports 2017, 7, 12970.

(45) Trowbridge, I. S.; Collawn, J. F.; Hopkins, C. R. Signal-Dependent Membrane Protein Trafficking in the Endocytic Pathway. Annu Rev Cell Biol 1993, 9, 129–161.

(46) Aguilar, R. C.; Wendland, B. Endocytosis of Membrane Receptors: Two Pathways Are Better Than One. Proceedings of the National Academy of Sciences 2005, 102, 2679–2680.

(47) Elkin, S. R.; Lakoduk, A. M.; Schmid, S. L. Endocytic Pathways and Endosomal Trafficking: A Primer. Wien Med Wochenschr 2016, 166, 196–204.

(48) Darwich, Z.; Klymchenko, A. S.; Dujardin, D.; Mély, Y. Imaging Lipid Order Changes in Endosome Membranes of Live Cells by Using a Nile Red-Based Membrane Probe. RSC Advances 2014, 4, 8481–8488.

(49) Kobayashi, T.; Beuchat, M. H.; Chevallier, J.; Makino, A.; Mayran, N.; Escola, J. M.; Lebrand, C.; Cosson, P.; Kobayashi, T.; Gruenberg, J. Separation and Characterization of Late Endosomal Membrane Domains. J Biol Chem 2002, 277, 32157–32164.

(50) Kolter, T.; Sandhoff, K. Lysosomal Degradation of Membrane Lipids. FEBS Lett 2010, 584, 1700–1712.

