## Supplemental information for publication. for "Covalent Solvatochromic Membrane Probe for Sensing Lipid Surrounding of Proteins"

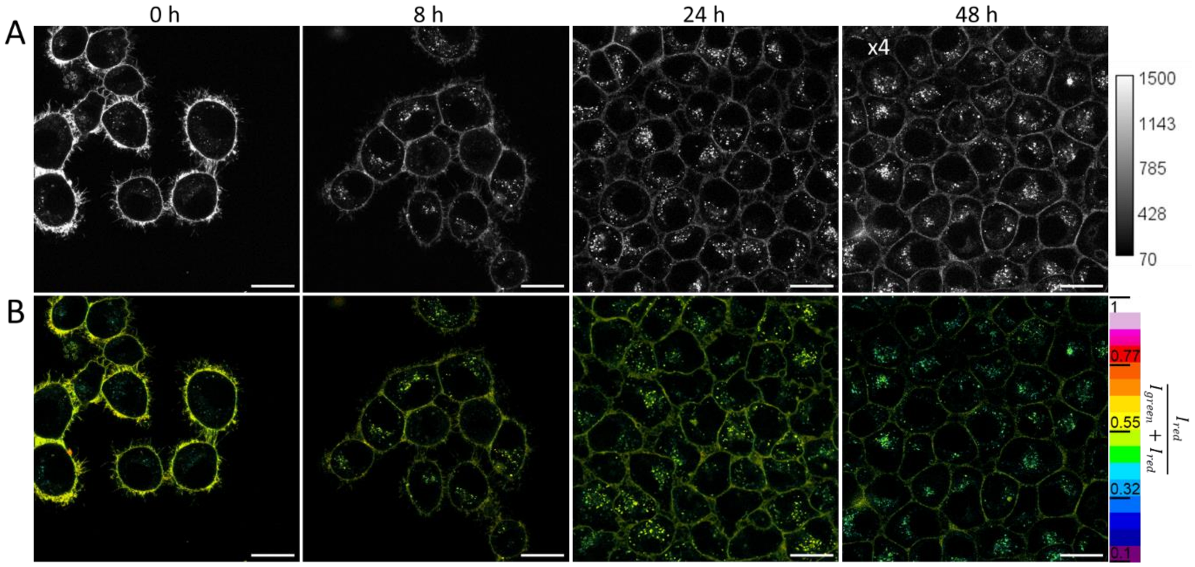


**Figure S1.** Confocal microscopy images of live KB cells stained with MemGraft-NR and left in full serum media with images taken at different time points. (A) intensity and (B) ratiometric images. Excitation wavelength was 488 nm. Scale bars: 20 µm.

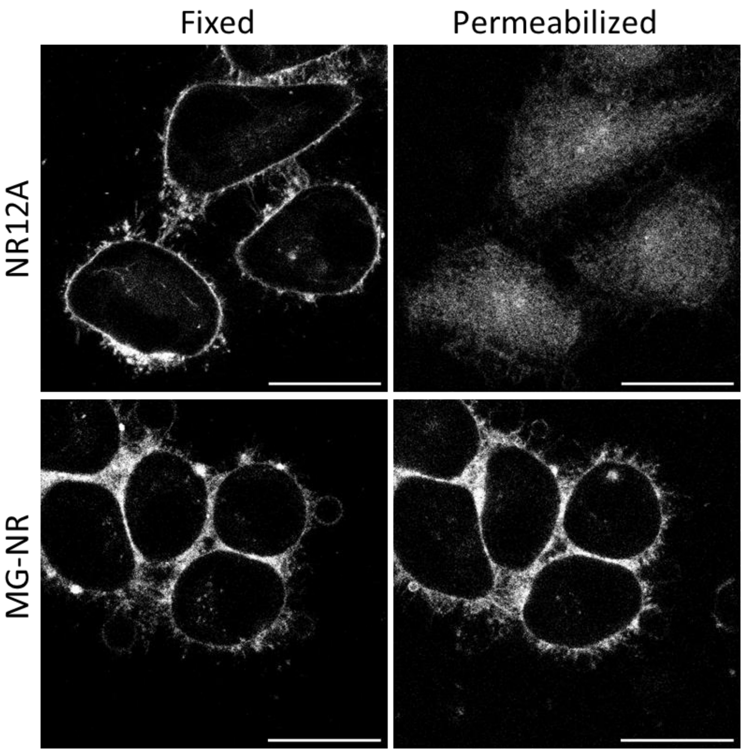


**Figure S2.** Confocal microscopy images KB cells stained with NR12A or MemGraft-NR after fixation (fixed) and the further permeabilization for 15 min in Tween 20 (permeabilized). Excitation wavelength was 488 nm. Scale bars: 20 µm.

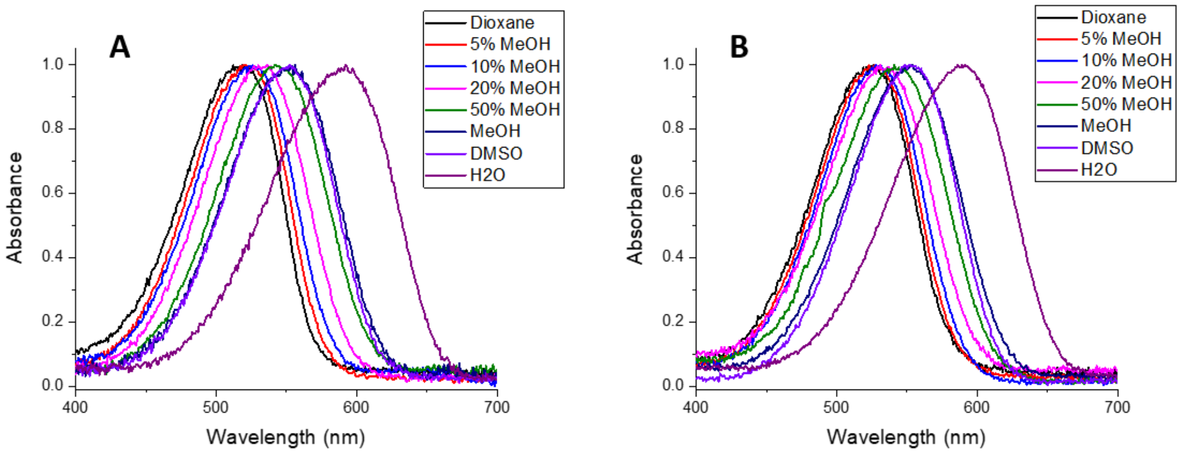


**Figure S3.** Normalized absorption spectra of Nile Red (A) and MG-NR (B)in solvents of different polarity and their mixtures. Dye concentration was 1 µM. Percentage of methanol indicates the amount of methanol in dioxane.

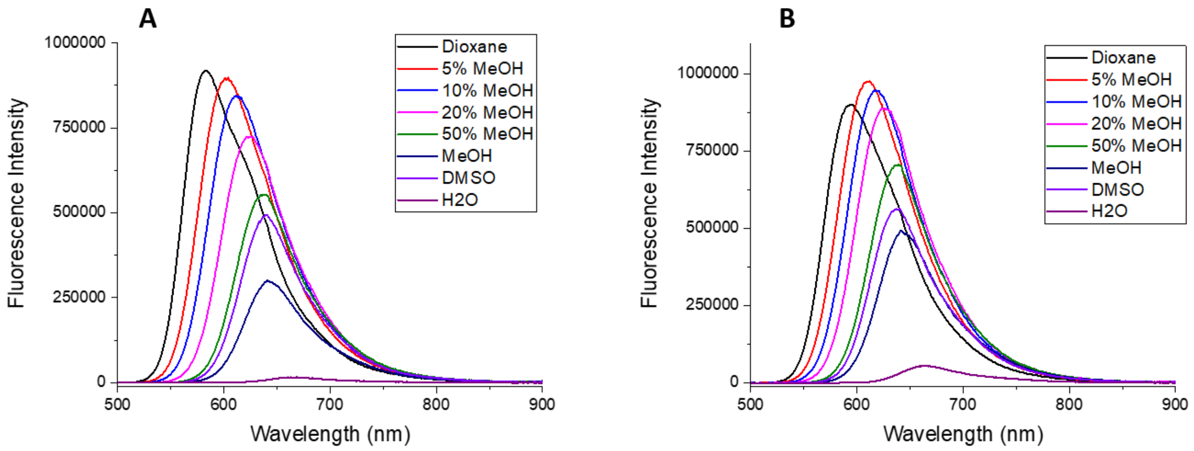


**Figure S4.** Fluorescence spectra of Nile Red (A) and MG-NR (B) in solvents of different polarity and their mixtures. Dye concentration was 1 µM. Excitation wavelength was 490 nm. Percentage of methanol indicates the amount of methanol in dioxane.

**Table S1.** Spectroscopic properties of Nile Red (NR) and MemGraft-NR (MG-NR) in solvents of different polarity and their mixtures. Dye concentration was 1 µM. Excitation wavelength was 490 nm.

| Solvent | Abs max NR | Abs max MG-NR | Fluo max NR | Fluo max MG-NR | QY NR | QY MG-NR |
|---|---|---|---|---|---|---|
| Dioxane | 519 | 526 | 584 | 595 | 0.70 (reference) | 0.66 |
| Diox + 5% MeOH | 520 | 523 | 604 | 612 | 0.75 | 0.75 |
| Diox + 10% MeOH | 526 | 528 | 611 | 616 | 0.76 | 0.73 |
| Diox + 20% MeOH | 536 | 531 | 621 | 625 | 0.68 | 0.74 |
| Diox + 50% MeOH | 542 | 541 | 637 | 639 | 0.65 | 0.60 |
| MeOH | 556 | 554 | 641 | 641 | 0.43 | 0.50 |
| DMSO | 552 | 552 | 639 | 640 | 0.87 | 0.82 |
| $H_2O$ | 592 | 591 | 669 | 666 | 0.06 | 0.12 |

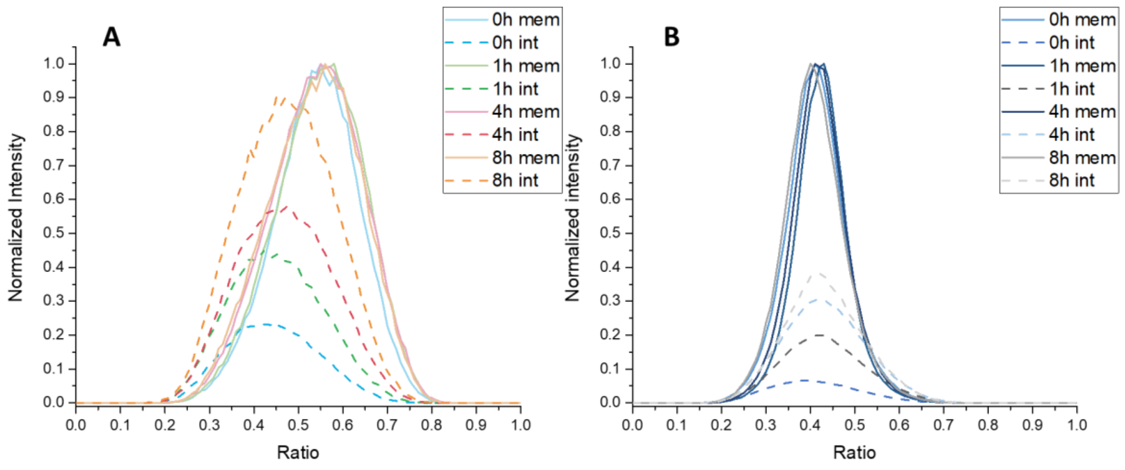


**Figure S5.** Ratiometric analysis of internalization of MemGraft-NR (A) and NR12A (B). The KB cell were incubated with the corresponding probe followed by two-color imaging and ratiometric analysis. The signals from the plasma membrane (solid line) and the cell interior (dashed curve) were separately analyzed. The relative differences in intensity between the PM and cell interior were respected during normalization for each timeframe.

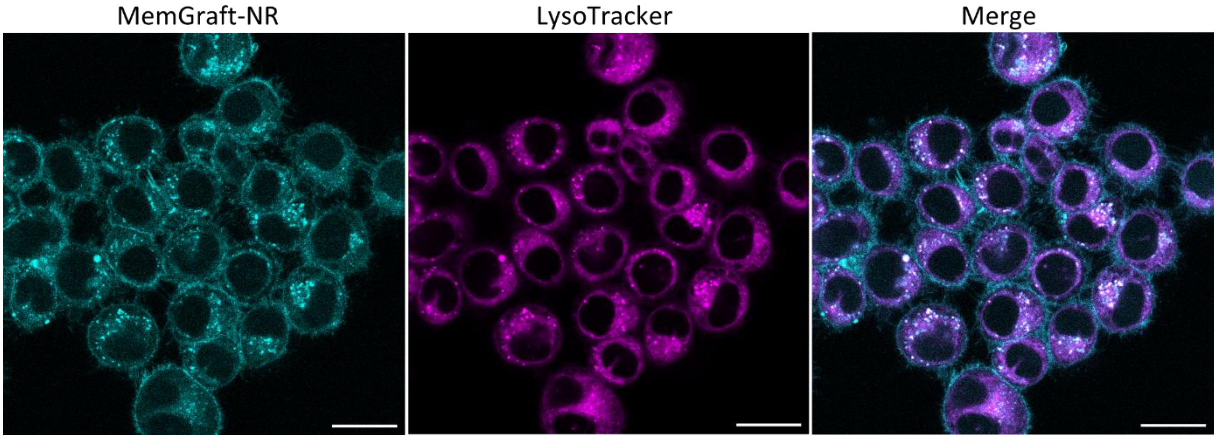


**Figure S6.** Confocal microscopy images of live KB cells in OptiMem stained with MemGraft-NR, then 3h incubation at 37 °C, followed by staining with LysoTracker Deep Red for 30 min at 37 °C. Scale bars: 20 µm.
